# A Cross-Sectional Study of Set Shifting Impairments and Falling in Individuals with and without Parkinson’s Disease

**DOI:** 10.1101/146332

**Authors:** J. Lucas McKay, Kimberly C. Lang, Lena H. Ting, Madeleine E. Hackney

## Abstract

INTRODUCTION. Individuals with Parkinson’s disease (PD) are at increased risk for falls, and exhibit deficits in executive function, including Set Shifting, which can be measured as the difference between parts B and A of the Trailmaking Test. METHODS. We conducted a cross-sectional study using baseline data of PD patients with and without freezing of gait (FOG) (n=69) and community-dwelling neurologically-normal older adults (NON-PD) (n=84) who had volunteered to participate in clinical rehabilitation research. Multivariate logistic regression analyses were performed to determine associations between Set Shifting, PD, and faller status, as determined by ≥1 self-reported falls in the previous 6 months, after adjusting for demographic and cognitive factors and clinical disease characteristics. RESULTS. Impaired Set Shifting was associated with previous falls after controlling for age, sex, overall cognitive function, PD, FOG, and PD disease duration (OR=1.29 [1.03-1.60]; P=0.02). In models controlling for age, sex, and overall cognitive function, PD was associated with increased fall prevalence among the study sample (OR=4.15 [95% CI 1.65-10.44], P<0.01) and FOG was associated with increased fall prevalence among the PD sample (OR=3.63 [1.22-10.80], P=0.02). Although the strongest associations between Set Shifting and falling were observed among PD without FOG (OR=2.11) compared to HOA (OR=1.14) and PD with FOG (OR=1.46) in a multivariate model that allowed for interaction between set shifting and PD status, there was insufficient evidence to reject the null hypothesis of no interaction. CONCLUSIONS. Set Shifting is associated with previous falls in non-demented older adults with and without PD.

**Highlights:**

- Individuals with PD are at increased risk for falls, although causes are unclear.
- Impaired Set Shifting was associated with falls in older adults with and without PD.
- Associations were strongest among those with PD but without freezing of gait.

## Introduction

Falls are a leading cause of accidental death (1), and fall risk is increased by about six times in individuals with Parkinson’s disease (PD) (2). Despite the significant morbidity and mortality resulting from falls, they remain extremely difficult to prevent due to their multifactorial causes (3). One of the strongest risk factors for falling among those with (4) and without PD (5) remains the presence of previous falls, which is of limited clinical utility for directing patients to interventions. Prospective studies have identified multiple disease-specific risk factors for falls among individuals with PD – including the presence of freezing of gait (FOG) – in addition to many of the generic or conventional fall risk factors identified in the aging population (3), but overall causes remain poorly understood.

Impaired executive function may play an important role in causing falls in individuals with and without PD (3, 6). Individuals with PD exhibit characteristic deficits in aspects of executive function, including Set Shifting (7, 8), a subdomain of executive function related to cognitive flexibility (9, 10). Set Shifting ability can be estimated as the difference between parts B and A of the Trailmaking Test (10, 11). PD is also associated with impaired Set Shifting in automatic motor responses during balance (12) and step initiation tasks (13), which suggests that impaired Set Shifting may cause impaired balance and falls in PD.

To the authors’ knowledge, no studies have attempted to relate impairments in the Set Shifting component of executive function to falling in individuals with or without PD. Here, we performed secondary analyses using existing baseline data of 153 adults with and without PD who had volunteered for exercise-based rehabilitation to test the hypotheses that: 1) impaired Set Shifting is associated with previous falls, and 2) that this association is modified by the presence of PD or PD and FOG.

## Methods

We assessed associations between impaired Set Shifting and previous falls using existing baseline measures of non-demented, community-dwelling individuals with and without PD from two previous exercise-based rehabilitative interventions designed to improve balance and mobility conducted in 2011-2013 and 2014-2015. Participants provided written informed consent according to protocols approved by the Institutional Review Boards of Emory University and the Georgia Institute of Technology. Participants met the following inclusion criteria: no diagnosed neurological conditions other than PD, ability to walk ≥3 meters with or without assistance. Participants with PD met the following additional inclusion criteria: diagnosis of idiopathic “definite PD” (14). Exclusion criteria were: significant musculoskeletal impairment as determined by the investigators.

Essential details of the rehabilitative intervention and outcome measures have been published previously (15-19). Briefly, participants were interviewed for health history and previous falls and assessed with a battery of behavioral and cognitive outcome measures prior to allocation to intervention arms with Adapted Tango rehabilitative dance classes or to control arms comprised of either standard care or health education classes.

Beginning with n=153 data records initially available for the present analysis, participants were excluded due to: presence of neurological conditions other than PD discovered after data collection (n=2), Montreal Cognitive Assessment (MoCA, (20)) scores (<18) indicating dementia (n=11), suspected invalid estimates of Set Shifting due to abnormally long times for Part A of the Trailmaking test (>200 seconds; n=2), and suspected invalid estimates of Set Shifting due to significant tremor artifacts in paper records of the Trailmaking test (n=1). After applying exclusions, there were data from n=138 individuals available for analysis.

### Study Variables

#### Primary outcome: Faller Status

The primary outcome was faller status. Participants were classified as “fallers” if they reported ≥1 falls (defined as “an event which results in a person coming to rest unintentionally on the ground or other lower level” (21)) in the prior six months at study entry.

#### Primary exposure: Set Shifting Score

The primary exposure, Set Shifting Score, was measured as the difference between Parts A and B of the Trailmaking Test. This timed test is administered on paper and requires the participant to quickly connect sequentially numbered dots (part A), or dots alternating between sequential numbers and letters (part B), including time required to correct any errors. Numerical scores for each part were truncated to 300 s and the difference between parts B and A was used as an estimate of Set Shifting impairment (10, 11). A larger difference indicates greater impairment in Set Shifting.

#### Secondary exposure: PD Status

The secondary exposure, PD Status, was treated as a dichotomous variable (NON-PD vs. PD, with NON-PD as the reference group) in univariate tests of central tendency, and as a trichotomous variable (NON-PD, PD-FOG, PD+FOG, with NON-PD as the reference group) in multivariate analyses. Participants with PD were classified as PD+FOG if they scored > 1 on item 3 of the Freezing of Gait Questionnaire (FOGQ) (22), indicating freezing more than once per week (15), and were classified as PD-FOG otherwise. Participants (n=5) for which FOGQ score was unavailable were classified as PD+FOG if they scored > 1 on item 14 of the Unified Parkinson’s Disease Rating Scale (UPDRS) Part II (23), indicating ‘occasional’ freezing (24).

Global cognitive function was assessed with the MoCA (20). PD disease severity was assessed with the UPDRS-III (23) by a Movement Disorders Society-trained examiner or by trained research assistants. Additional study variables considered to be relevant for evaluating associations with falling included the demographic and clinical variables moderately or significantly associated with elevated fall risk in PD, including age, female sex, and selfreported PD duration in years (4). Additional motor domain variables included Berg Balance Scale (BBS) (3, 25) and self-selected gait speed (4, 26). MoCA score was dichotomized about 27, with scores ≤26 indicating mild cognitive impairment (MCI; mocatest.org). BBS score was dichotomized about 45, indicating functional mobility without the use of a cane (25), and gait speed was dichotomized about 0.7 m/s, a previously-reported cutoff for slow gait (26).

### Analytic Plan

#### Descriptive statistics and univariate tests of central tendency across groups

Descriptive statistics were calculated for study variables overall and stratified on PD status. Imbalances across groups were assessed with univariate tests of central tendency (independent sample *t*-tests, Wilcoxon rank sum, chi-square) between the NON-PD and PD strata, and between the PD-FOG and PD+FOG strata within the PD group. Satterthwaite’s formula was applied to calculate variances as necessary when equal variance assumptions were unreasonable based on the Folded F statistic. In cases where the total sample size was <40, exact Wilcoxon rank sum tests were performed. Exact Wilcoxon rank sum tests were also performed for Parts A and B of the Trailmaking Test, and for Set Shifting Score due to the strong right tail observed in the distribution of these variables. Differences between groups in proportions were assessed with two-tailed chi-squared tests.

#### Multivariate associations between Set Shifting Score and Faller Status

Multivariate logistic regression models were used to estimate associations between Set Shifting Score, PD Status, and the primary outcome Faller Status. Associations were expressed as prevalence odds ratios (OR) ±95% confidence intervals (CI). Set Shifting Score was expressed with respect to the minimum value observed in the sample and scaled to units of 30 seconds, approximately one quartile. Odds ratios were calculated in unadjusted models and in models adjusted for sex, age (in 5-year units), presence of MCI, and PD duration (in 5-year units).

To test whether Set Shifting score was associated with previous falls, we fit the following multivariate model:

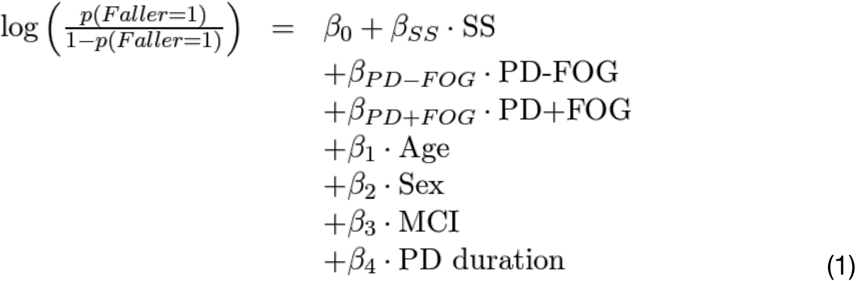

where variable SS indicates Set Shifting Score, indicator variable PD-FOG is 1 for individuals in the PD-FOG stratum and 0 otherwise, and indicator variable PD+FOG is 1 for individuals in the PD+FOG stratum and 0 otherwise. To test whether impaired Set Shifting was associated with previous falls, the following null hypothesis was evaluated with a Wald test:

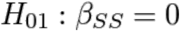

To test whether the association between Set Shifting and previous falls was modified by the presence of PD or PD and FOG, the parameters of a second adjusted multivariate model allowing interaction between Set Shifting Score and PD Status were also estimated:

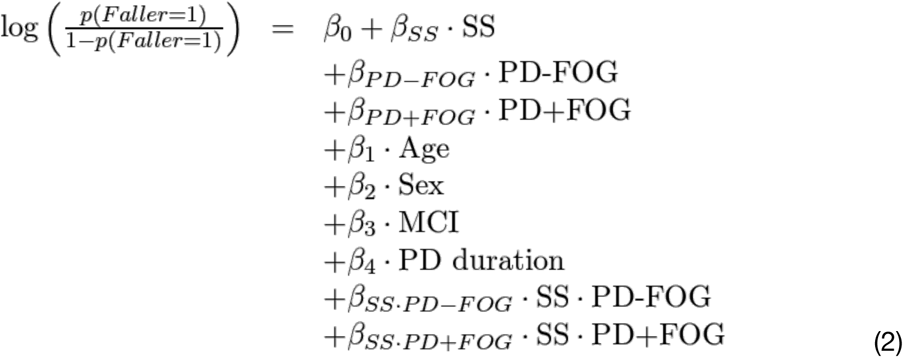

A likelihood ratio test was then employed comparing the full model (Eq. 2) against the reduced model (Eq. 1) to evaluate the following null hypothesis:

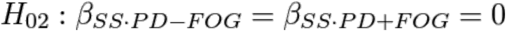

#### Additional analyses

To minimize the potential for misclassification bias associated with retrospective self-report of previous falls, results of the adjusted model (Eq. 1) were compared after imposing a more stringent criteria for faller status. In this analysis, participants were classified as “fallers” if they reported ≥2 falls in the previous 6 months. Sensitivity of the adjusted model (Eq. 1) to the inclusion of motor domain covariates BBS and gait speed was also assessed. Finally, to facilitate comparisons with prior studies, additional multivariate logistic regression models were also calculated to estimate prevalence odds ratios for PD vs. NON-PD and for PD+FOG vs. PD-FOG with Set Shifting Score omitted.

Due to the exploratory nature of the study no a priori power analyses were performed. All reported P-values correspond to 2-tailed tests considered statistically-significant at P<0.05. Analyses were performed using SAS University Edition.

## Results

Demographic and clinical characteristics of the study population stratified on the presence of PD and on the presence of FOG are presented in Tables 1 and 2. Overall prevalence of previous falls was 51/138=40%. Participants with PD exhibited significantly increased fall prevalence (34/65=52% vs. 17/73=23%, P<0.01) despite being younger, higher functioning cognitively, and less likely to be female than the NON-PD group, all of which are known fall risk factors (3). Among the PD group, individuals with and without FOG were relatively well-matched on demographic variables, cognitive function, and disease duration (Table 2); FOG was associated with more severe UPDRS-III score, poorer BBS score, more impaired Set Shifting, and increased prevalence of previous falls (18/26=69% vs. 16/39=40%).

**Table 1.**
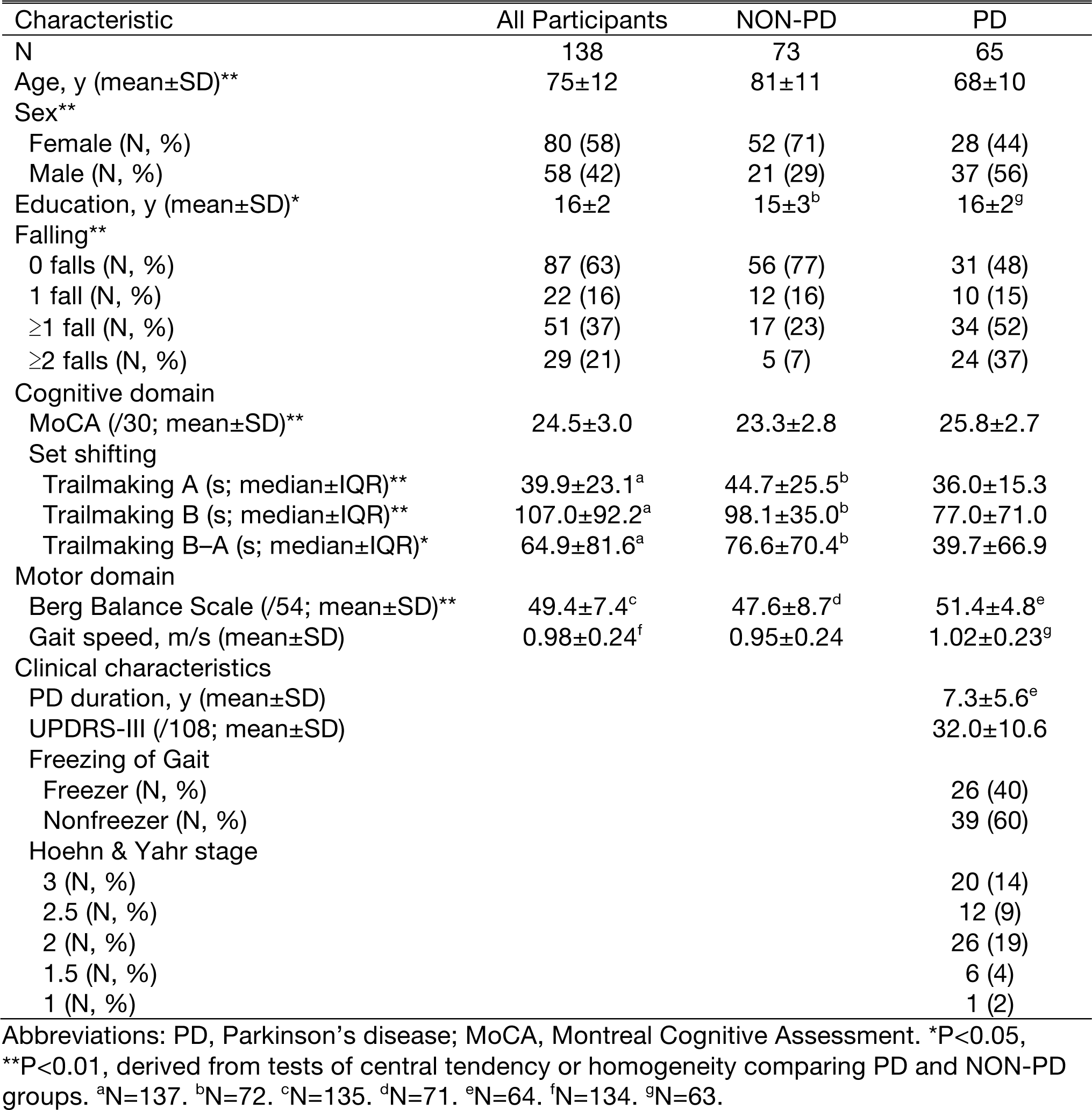
Demographic and clinical features of the study population, assembled from baseline measurements of rehabilitative interventions conducted in 2011-2013 and 2014-2015, overall

| Characteristic | All Participants | NON-PD | PD |
| --- | --- | --- | --- |
| N | 138 | 73 | 65 |
| Age, y (mean±SD)** | 75±12 | 81±11 | 68±10 |
| Sex** |  |  |  |
| Female (N, %) | 80 (58) | 52 (71) | 28 (44) |
| Male (N, %) | 58 (42) | 21 (29) | 37 (56) |
| Education, y (mean±SD)* | 16±2 | 15±3<sup>b</sup> | 16±2<sup>g</sup> |
| Falling** |  |  |  |
| 0 falls (N, %) | 87 (63) | 56 (77) | 31 (48) |
| 1 fall (N, %) | 22 (16) | 12 (16) | 10 (15) |
| ≥1 fall (N, %) | 51 (37) | 17 (23) | 34 (52) |
| ≥2 falls (N, %) | 29 (21) | 5 (7) | 24 (37) |
| Cognitive domain |  |  |  |
| MoCA (/30; mean±SD)** | 24.5±3.0 | 23.3±2.8 | 25.8±2.7 |
| Set shifting |  |  |  |
| Trailmaking A (s; median±IQR)** | 39.9±23.1<sup>a</sup> | 44.7±25.5<sup>b</sup> | 36.0±15.3 |
| Trailmaking B (s; median±IQR)** | 107.0±92.2<sup>a</sup> | 98.1±35.0<sup>b</sup> | 77.0±71.0 |
| Trailmaking B–A (s; median±IQR)* | 64.9±81.6<sup>a</sup> | 76.6±70.4<sup>b</sup> | 39.7±66.9 |
| Motor domain |  |  |  |
| Berg Balance Scale (/54; mean±SD)** | 49.4±7.4<sup>c</sup> | 47.6±8.7<sup>d</sup> | 51.4±4.8<sup>e</sup> |
| Gait speed, m/s (mean±SD) | 0.98±0.24<sup>f</sup> | 0.95±0.24 | 1.02±0.23<sup>g</sup> |
| Clinical characteristics |  |  |  |
| PD duration, y (mean±SD) |  |  | 7.3±5.6<sup>e</sup> |
| UPDRS-III (/108; mean±SD) |  |  | 32.0±10.6 |
| Freezing of Gait |  |  |  |
| Freezer (N, %) |  |  | 26 (40) |
| Nonfreezer (N, %) |  |  | 39 (60) |
| Hoehn & Yahr stage |  |  |  |
| 3 (N, %) |  |  | 20 (14) |
| 2.5 (N, %) |  |  | 12 (9) |
| 2 (N, %) |  |  | 26 (19) |
| 1.5 (N, %) |  |  | 6 (4) |
| 1 (N, %) |  |  | 1 (2) |

Abbreviations: PD, Parkinson's disease; MoCA, Montreal Cognitive Assessment. \*P<0.05, \*\*P<0.01, derived from tests of central tendency or homogeneity comparing PD and NON-PD groups. <sup>a</sup>N=137. <sup>b</sup>N=72. <sup>c</sup>N=135. <sup>d</sup>N=71. <sup>e</sup>N=64. <sup>f</sup>N=134. <sup>g</sup>N=63.

**Table 2.** Demographic and clinical features of PD patients in the study population, assembled from baseline measurements of rehabilitative interventions conducted in 2011-2013 and 2014-2015, stratified on the presence of freezing of gait (FOG).

| Characteristic | PD-FOG | PD+FOG |
| --- | --- | --- |
| N | 39 | 26 |
| Age, y (mean±SD) | 69±8 | 67±12 |
| Sex |  |  |
| Female (N, %) | 18 (46) | 10 (38) |
| Male (N, %) | 21 (54) | 16 (62) |
| Education, y (mean±SD) | 16±2<sup>c</sup> | 16±2<sup>a</sup> |
| Falling* |  |  |
| 0 falls (N, %) | 23 (60) | 8 (31) |
| 1 fall (N, %) | 8 (20) | 2 (8) |
| ≥1 fall (N, %) | 16 (40) | 18 (69) |
| ≥2 falls (N, %) | 8 (20) | 16 (61) |
| Cognitive domain |  |  |
| MoCA (/30; mean±SD) | 26.1±2.7 | 25.4±2.6 |
| Set shifting |  |  |
| Trailmaking A (s; median±IQR)* | 29.8±11.0 | 41.1±16.3<sup>a</sup> |
| Trailmaking B (s; median±IQR)* | 65.5±54.7 | 113.1±88.6<sup>a</sup> |
| Trailmaking B–A (s; median±IQR)* | 34.3±44.2 | 69.8±71.6<sup>a</sup> |
| Motor domain |  |  |
| Berg Balance Scale (/54; mean±SD)* | 52.9±3.3 | 49.2±6.0 |
| Gait speed, m/s (mean±SD) | 1.06±0.21 | 0.95±0.26<sup>b</sup> |
| Clinical characteristics |  |  |
| PD duration, y (mean±SD) | 6.4±5.8 | 8.5±5.1<sup>a</sup> |
| UPDRS-III (/108; mean±SD)* | 29.4±7.8 | 35.9±13.0 |
| Freezing of Gait |  |  |
| Freezer (N, %) | 0 (0) | 26 (100) |
| Nonfreezer (N, %) | 39 (100) | 0 (0) |
| Hoehn & Yahr stage |  |  |
| 3 (N, %) | 9 (23) | 11 (42) |
| 2.5 (N, %) | 6 (15) | 6 (23) |
| 2 (N, %) | 17 (43) | 9 (35) |
| 1.5 (N, %) | 6 (15) | 0 (0) |
| 1 (N, %) | 1 (3) | 0 (0) |

Abbreviations: PD, Parkinson's disease; MoCA, Montreal Cognitive Assessment. \*P<0.05, derived from tests of central tendency or homogeneity comparing PD+FOG and PD-FOG groups. <sup>a</sup>N=25. <sup>b</sup>N=24. <sup>c</sup>N=38.

### Associations with previous falls

Model (Eq. 1) demonstrated a significant association between impaired Set Shifting and previous falls (OR: 1.29, 95% CI: 1.03, 1.60; P<0.02) after adjusting for age, sex, PD duration, and presence of MCI. PD Status was also significantly associated with previous falls (PD+FOG OR: 4.69, 95% CI: 1.30, 16.98; P<0.02); however, contrasts between the PD+FOG and PD-FOG groups (OR: 1.64) were not statistically significant. Comparable associations between Set Shifting and previous falls were observed in a model that was unadjusted for age, sex, PD duration, and presence of MCI (OR: 1.19, 95% CI: 0.99, 1.44); however, associations were statistically significant only in the adjusted model (Table 3).

**Table 3.** Associations between Set Shifting Score, PD Status, and ≥1 falls in the previous 6 months in the study sample (Model 1).

|  | Unadjusted |  |  | Adjusted<sup>a</sup> |  |  |
| --- | --- | --- | --- | --- | --- | --- |
|  | OR | 95% CI | P | OR | 95% CI | P |
| Set Shifting Score | 1.19 | 0.99, 1.44 | 0.07 | 1.29 | 1.03, 1.60 | 0.02 |
| PD-FOG vs. NON-PD | 2.90 | 1.18, 7.14 | 0.02 | 2.87 | 0.92, 8.90 | 0.07 |
| PD+FOG vs. NON-PD | 7.50 | 2.68, 21.00 | <0.01 | 4.69 | 1.30, 16.98 | 0.02 |
| PD+FOG vs. PD-FOG | 2.59 | 0.88, 7.62 | <0.01 | 1.64 | 0.46, 5.84 | 0.45 |
| No. Obs |  | 136 |  |  | 135 |  |
| No. Events |  | 50 |  |  | 49 |  |
| -2•Ln(L) |  | 159.348 |  |  | 137.897 |  |

Abbreviations: PD, Parkinson's disease; FOG, freezing of gait; OR, odds ratio; CI, confidence interval. <sup>a</sup>Adjusted for age, sex, PD duration, MCI.

Likelihood ratio tests comparing Model (Eq. 2), which allowed interaction between Set Shifting Score and PD Status, to Model (Eq. 1) demonstrated that the association between Set Shifting and previous falls did not vary in a statistically significant fashion across the NON-PD, PD-FOG, and PD+FOG groups. Results were comparable with (P-interaction=0.21) or without (P-interaction=0.34) adjustments for age, sex, PD duration, and MCI. Although not statistically significant, qualitatively stronger associations between Set Shifting and previous falls were observed among the PD-FOG group (adjusted OR=2.11, 95% CI: 0.94, 4.70) compared to either among the NON-PD group (OR=1.14, 95% CI: 0.86, 1.50) or among the PD+FOG group (OR=1.46, 95% CI: 0.96, 2.23) (Table 4).

**Table 4.**
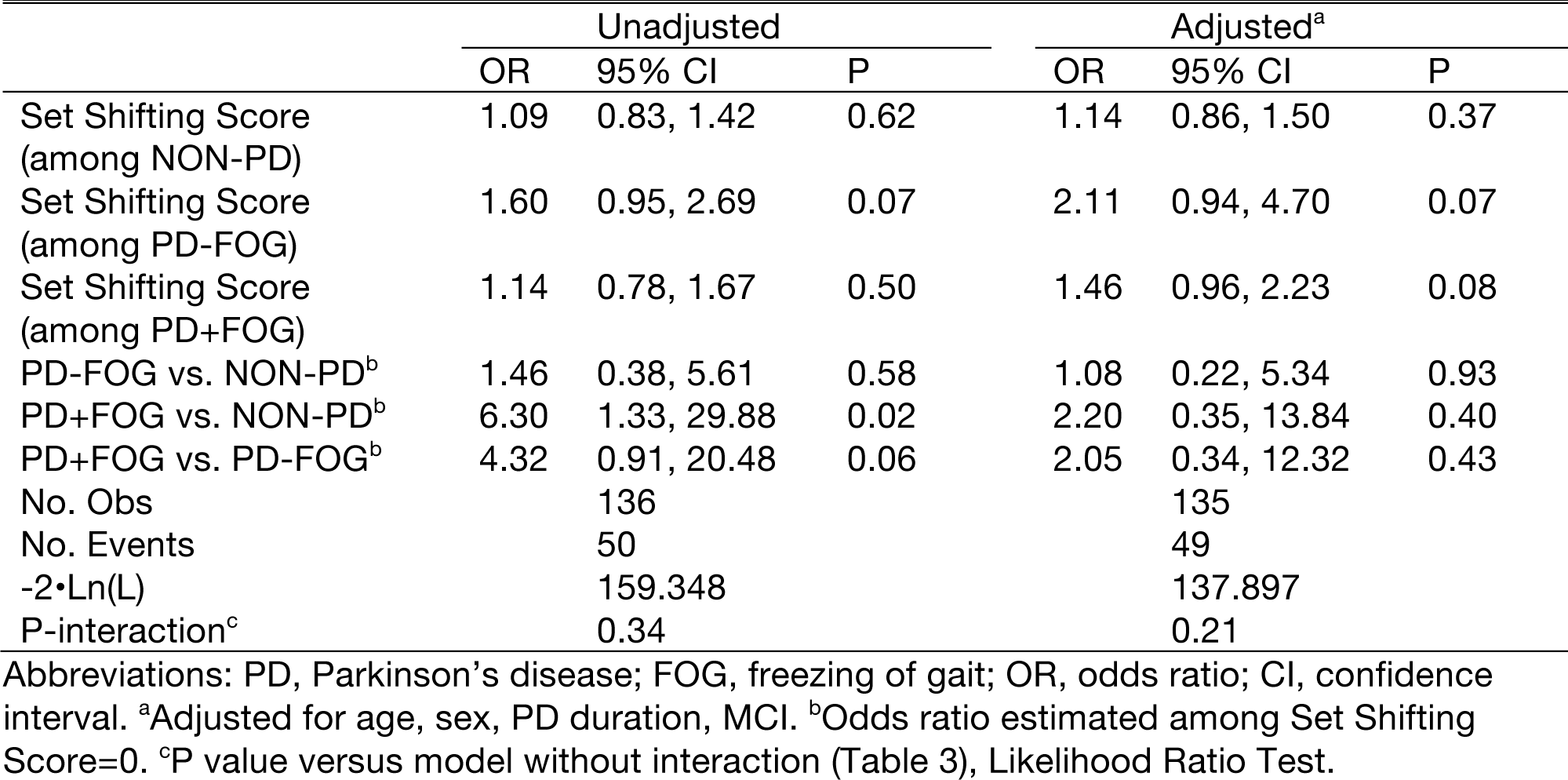
Associations between Set Shifting Score, PD Status, and ≥1 falls in the previous 6 months in the study sample, allowing for interaction between Set Shifting Score and PD Status (Model 2).

### Additional analyses

Associations between Set Shifting and previous falls were essentially unchanged when a more stringent definition of faller status was imposed (Table S1; OR: 1.28 vs. 1.29 in adjusted Model 1). Unlike the main model, contrasts between PD+FOG and PD-FOG were statistically significant (OR: 4.28, CI: 1.14, 16.16, P<0.03) under a more stringent definition of faller status. Including motor domain covariates in the model affected identified odds ratios by ≈10%, reducing odds ratios for Set Shifting (OR: 1.21, vs. 1.29) and PD-FOG (2.66 vs. 2.87) and increasing odds ratios for PD+FOG (5.06 vs. 4.69) (Table S2). In multivariate models controlling for age, sex, and MCI, but without Set Shifting, odds ratio contrasting PD to NON-PD was 4.15 (CI: 1.65, 10.44) and the odds ratio contrasting PD+FOG to PD-FOG was 3.63 (CI: 1.22, 10.80).

## Discussion

Consistent with our hypothesis, in this cross-sectional study of 138 non-demented individuals with and without PD, impaired Set Shifting was associated with previous falls after controlling for demographic and clinical variables as well as for overall cognitive function.

Consistent with prospective studies in the literature, we also identified very strong associations between disease state and previous falls, providing confidence in these results. In models adjusted for age, sex, and MCI, the odds of previous falls were elevated >4 times among those with PD compared to those without, which largely agrees with other work. Similarly, among those with PD, the odds of previous falls were elevated >3 times among those with FOG. Recent prospective studies have identified generally comparable odds ratios (PD OR: 6.08, CI: 2.45, 15.05 (2); PD+FOG OR: 4.11, CI: 2.20, 7.66 (4)). It is unknown why the odds ratios identified here were biased downward somewhat compared to values from the literature. Results were essentially unchanged under a more stringent definition of “faller,” suggesting that this bias was not due to the use of self-reported fall history. We speculate that these biases may result from elevated fall prevalence among the NON-PD group, some of whom might have enrolled in the rehabilitative program due to concerns about previous falls.

The association between Set Shifting score and previous falls observed here supports the hypothesis that impairments in subdomains of executive function – rather than overall cognitive function – may be associated with falls in individuals with and without PD. Possibly this relationship may be observed because impaired Set Shifting makes motor tasks more challenging. Other measures of executive function have been associated with increased fall risk in non-demented PD patients (27, 28) and in neurologically-intact older adults (6). Causal links between impaired Set Shifting and falling are unclear, but at least among PD patients, impaired Set Shifting during motor domain tasks such as reactive balance (12) and step initiation (13) may provide a possible causal pathway between impaired Set Shifting and falling.

Inconsistent with our hypothesis, we found only qualitative evidence that associations between Set Shifting and falls were modified by disease state, which casts doubt on the hypothesis that PD-specific (13) or FOG-specific (10) impairments in Set Shifting, at least, are associated with falls. Qualitatively, the strongest associations between Set Shifting and previous falls were observed in PD-FOG (OR 2.11). This suggests that people with PD but without FOG could benefit from pharmacological or training-based interventions aimed at improving cognitive function and mitigating fall risk. However, we could not reject the null hypothesis that the association was constant across study strata. This important question could be addressed in a larger, prospective study.

This study has some limitations of note. First, although motor domain variables have been demonstrated to predict incident falls in prospective studies (4, 26), we were unable to control for these variables in the main models of this cross-sectional study because of the potential for reversed causality. Specifically, we could not eliminate the possibility that impairments in these variables could have resulted from – rather than contributed to – previous falls (29). Therefore, sensitivity to these variables was assessed, but this model was not used for primary hypothesis tests. Further, although we attempted to minimize misclassification error associated with self-report of FOG status by using a robust classification for FOG, this process was likely imperfect and may have reduced power to discriminate between groups.

In summary, impaired Set Shifting was associated with previous falls in non-demented individuals with and without PD. The strongest associations were observed among individuals with PD but without FOG, although there was insufficient evidence to distinguish this interaction effect from the null.

## Author roles

Research project: Conception, JLM, LHT, MEH; Organization, JLM, KCL, LHT, MEH; Execution, JLM, KCL, LHT, MEH. Statistical Analysis: Design and Execution: JLM; Review and Critique: LHT, MEH. Manuscript Preparation: Writing of the first draft: JLM; Review and Critique, KCL, LHT, MEH.

## Supplementary Information

**Table S1.** Associations between Set Shifting Score, PD Status, and ≥ 2 falls in the previous 6 months in the study sample.

|  | Adjusted<sup>a</sup> |  |  |
| --- | --- | --- | --- |
|  | OR | 95% CI | P |
| Set Shifting Score | 1.28 | 0.99, 1.66 | 0.06 |
| PD-FOG vs. NON-PD | 4.10 | 0.90, 18.69 | 0.07 |
| PD+FOG vs. NON-PD | 17.54 | 3.76, 81.85 | <0.01 |
| PD+FOG vs. PD-FOG | 4.28 | 1.14, 16.16 | 0.03 |
| No. Obs |  | 135 |  |
| No. Events |  | 27 |  |
| -2•Ln(L) |  | 92.251 |  |

Abbreviations: PD, Parkinson's disease; FOG, freezing of gait; OR, odds ratio; CI, confidence interval. <sup>a</sup>Adjusted for age, sex, PD duration, MCI.

**Table S2.** Associations between Set Shifting Score, PD Status, and ≥1 falls in the previous 6 months in the study sample, adjusted for motor domain covariates.

|  | Further Adjusted<sup>a</sup> |  |  |
| --- | --- | --- | --- |
|  | OR | 95% CI | P |
| Set Shifting Score | 1.21 | 0.95, 1.53 | 0.12 |
| PD-FOG vs. NON-PD | 2.66 | 0.82, 8.60 | 0.10 |
| PD+FOG vs. NON-PD | 5.06 | 1.29, 19.87 | 0.02 |
| PD+FOG vs. PD-FOG | 1.91 | 0.49, 7.43 | 0.35 |
| No. Obs |  | 130 |  |
| No. Events |  | 48 |  |
| -2•Ln(L) |  | 129.087 |  |

Abbreviations: PD, Parkinson's disease; FOG, freezing of gait; OR, odds ratio; CI, confidence interval. <sup>a</sup>Further adjusted for age, sex, PD duration, mild cognitive impairment, Berg Balance Scale, and self-selected gait speed.

## References

1. Deandrea S, Lucenteforte E, Bravi F, Foschi R, La Vecchia C, Negri E. Risk factors for falls in community-dwelling older people: a systematic review and meta-analysis. Epidemiology. 2010;21(5):658–68. doi: 10.1097/EDE.0b013e3181e89905

2. Bloem BR, Grimbergen YAM, Cramer M, Willemsen M, Zwinderman AH. Prospective assessment of falls in Parkinson’s disease. J Neurol. 2001;248(11):950–8. doi: 10.1007/s004150170047

3. van der Marck MA, Klok MP, Okun MS, Giladi N, Munneke M, Bloem BR, et al. Consensus-based clinical practice recommendations for the examination and management of falls in patients with Parkinson’s disease. Parkinsonism Relat Disord. 2014;20(4):360–9. doi: 10.1016/j.parkreldis.2013.10.030

4. Paul SS, Canning CG, Sherrington C, Lord SR, Close JC, Fung VS. Three simple clinical tests to accurately predict falls in people with Parkinson’s disease. Mov Disord. 2013;28(5):655–62. doi: 10.1002/mds.25404

5. Tinetti ME, Speechley M, Ginter SF. Risk factors for falls among elderly persons living in the community. The New England journal of medicine. 1988;319(26):1701–7. doi: 10.1056/NEJM198812293192604

6. Mirelman A, Herman T, Brozgol M, Dorfman M, Sprecher E, Schweiger A, et al. Executive function and falls in older adults: new findings from a five-year prospective study link fall risk to cognition. PloS one. 2012;7(6):e40297. doi: 10.1371/journal.pone.0040297

7. Woodward TS, Bub DN, Hunter MA. Task switching deficits associated with Parkinson’s disease reflect depleted attentional resources. Neuropsychologia. 2002;40(12):1948–55. doi:

8. Dirnberger G, Jahanshahi M. Executive dysfunction in Parkinson’s disease: a review. J Neuropsychol. 2013;7(2):193–224. doi: 10.1111/jnp.12028

9. Miyake A, Friedman NP, Emerson MJ, Witzki AH, Howerter A, Wager TD. The unity and diversity of executive functions and their contributions to complex “Frontal Lobe” tasks: a latent variable analysis. Cogn Psychol. 2000;41(1):49–100. doi: 10.1006/cogp.1999.0734

10. Cohen RG, Klein KA, Nomura M, Fleming M, Mancini M, Giladi N et al. Inhibition, executive function and freezing of gait. Journal of Parkinson’s disease. 2014;4(1):111–22. doi: 10.3233/JPD-130221

11. Factor SA, Scullin MK, Sollinger AB, Land JO, Wood-Siverio C, Zanders L, et al. Freezing of gait subtypes have different cognitive correlates in Parkinson’s disease. Parkinsonism Relat Disord. 2014;20(12):1359–64. doi: 10.1016/j.parkreldis.2014.09.023

12. Dimitrova D, Horak FB, Nutt JG. Postural Muscle Responses to Multidirectional Translations in Patients With Parkinson’s Disease. J Neurophysiol. 2004;91(1):489–501. doi:

13. Smulders K, Esselink RA, Bloem BR, Cools R. Freezing of gait in Parkinson’s disease is related to impaired motor switching during stepping. Mov Disord. 2015;30(8):1090–7. doi: 10.1002/mds.26133

14. Racette BA, Rundle M, Parsian A, Perlmutter JS. Evaluation of a screening questionnaire for genetic studies of Parkinson’s disease. Am J Med Genet. 1999;88(5):539–43. doi:

15. McKee KE, Hackney ME. The effects of adapted tango on spatial cognition and disease severity in Parkinson’s disease. Journal of motor behavior. 2013;45(6):519–29. doi: 10.1080/00222895.2013.834288

16. Hackney ME, Byers C, Butler G, Sweeney M, Rossbach L, Bozzorg A. Adapted Tango Improves Mobility Motor-Cognitive Function and Gait but Not Cognition in Older Adults in Independent Living. Journal of the American Geriatrics Society. 2015;63(10):2105–13. doi: 10.1111/jgs.13650

17. McKay J, Ting L, Hackney M. Balance, Body Motion and Muscle Activity After High-Volume Short-Term Dance-Based Rehabilitation in Persons With Parkinson Disease: A Pilot Study. J Neurol Phys Ther. 2016;40(4):257–68. doi: 10.1097/NPT.0000000000000150

18. Hackney ME, Earhart GM. Effects of dance on gait and balance in Parkinson’s disease: a comparison of partnered and nonpartnered dance movement. Neurorehabilitation and neural repair. 2010;24(4):384–92. doi:

19. Hackney ME, Earhart GM. Recommendations for implementing partnered dance classes for persons with Parkinson Disease. Am J Dance Ther. 2010;31(1):41–5. doi:

20. Hoops S, Nazem S, Siderowf AD, Duda JE, Xie SX, Stern MB et al. Validity of the MoCA and MMSE in the detection of MCI and dementia in Parkinson disease. Neurology. 2009;73(21):1738–45. doi: 10.1212/WNL.0b013e3181c34b47

21. Ashburn A, Stack E, Pickering RM, Ward CD. Predicting fallers in a community-based sample of people with Parkinson’s disease. Gerontology. 2001;47(5):277–81. doi: 52812

22. Giladi N, Shabtai H, Simon ES, Biran S, Tal J, Korczyn AD. Construction of freezing of gait questionnaire for patients with Parkinsonism. Parkinsonism Related Disord. 2000;6(3):165–70. doi: 10.1016/s1353-8020(99)00062-0

23. Fahn S, Elton RL, Members of the UPDRS Development Committee. The Unified Parkinson’s Disease Rating Scale. In: Fahn S, Marsden CD, Calne DB, Goldstein M, editors. Recent Developments in Parkinson’s Disease. 2. Florham Park NJ: Macmillan Healthcare Information; 1987. p. 153–63.

24. Perez-Lloret S, Negre-Pages L, Damier P, Delval A, Derkinderen P, Destee A, et al. Prevalence, determinants, and effect on quality of life of freezing of gait in Parkinson disease. JAMA neurology. 2014;71(7):884–90. doi: 10.1001/jamaneurol.2014.753

25. Berg KO, Maki BE, Williams JI, Holliday PJ, Wood-Dauphinee SL. Clinical and laboratory measures of postural balance in an elderly population Archives of physical medicine and rehabilitation. 1992;73(11):1073–80. doi:

26. Verghese J, Holtzer R, Lipton RB, Wang C. Quantitative gait markers and incident fall risk in older adults. The journals of gerontology Series A Biological sciences and medical sciences. 2009;64(8):896–901. doi: 10.1093/gerona/glp033

27. Latt MD, Lord SR, Morris JG, Fung VS. Clinical and physiological assessments for elucidating falls risk in Parkinson’s disease. Mov Disord. 2009;24(9):1280–9. doi: 10.1002/mds.22561

28. Mak MK, Wong A, Pang MY. Impaired executive function can predict recurrent falls in Parkinson’s disease. Archives of physical medicine and rehabilitation. 2014;95(12):2390–5. doi: 10.1016/j.apmr.2014.08.006

29. Salzman B. Gait and balance disorders in older adults. Am Fam Physician. 2010;82(1):61–8. doi:

